# Proteomic Analysis of Endemic Viral Infections in Neurons offers Insights into Neurodegenerative Diseases

**DOI:** 10.1101/2025.03.17.643709

**Authors:** Ziyi Li, Negin P. Martin, Jacob Epstein, Shih-Heng Chen, Ying Hao, Daniel M. Ramos, Kate M. Andersh, Paige Jarreau, Cory Weller, Mike A. Nalls, Caroline B. Pantazis, Luigi Ferrucci, Mark R. Cookson, Andrew B. Singleton, Yue Andy Qi, Jerrel L. Yakel

## Abstract

Endemic viral infections with low pathogenicity are often overlooked due to their mild symptoms, yet they can exert long-term effects on cellular function and contribute to disease pathogenesis. While viral infections have been implicated in neurodegenerative disorders, their impact on the neuronal proteome remains poorly understood. Here, we differentiated human induced pluripotent stem cells (KOLF2.1J) into mature neurons to investigate virus-induced proteomic changes following infection with five neurotropic endemic human viruses: Herpes simplex virus 1 (HSV-1), Human coronavirus 229E (HCoV-229E), Epstein-Barr virus (EBV), Varicella-Zoster virus (VZV), and Influenza A virus (H1N1). Given that these viruses can infect adults and have the potential to cross the placental barrier, their molecular impact on neurons may be relevant across the lifespan. Using mass spectrometry-based proteomics with a customized library for simultaneous detection of human and viral proteins, we confirmed successful infections and identified virus-specific proteomic signatures. Notably, virus-induced protein expression changes converged on key neuronal pathways, including those associated with neurodegeneration. Gene co-expression network analysis identified protein modules correlated with viral proteins. Pathway enrichment analysis of these modules revealed associations with the nervous system, including pathways linked to Alzheimer’s and Parkinson’s disease. Remarkably, several viral-induced proteomic alterations overlapped with changes observed in postmortem Alzheimer’s patient brains, suggesting a mechanistic connection between viral exposure and neurodegenerative disease progression. These findings provide molecular insights into how common viral infections perturb neuronal homeostasis and may contribute to neurodegenerative pathology, highlighting the need to consider endemic viruses as potential environmental risk factors in neurological disorders.

## Introduction

Endemic viruses are highly adapted to human-to-human transmission, maintaining persistent reservoirs within populations despite the build-up of host immunity ^1^. These viruses typically circulate at low levels, ensuring their continued presence within the population. Their transmission patterns are often predictable, yet they can evolve mechanisms to evade immune defenses or persist in sanctuary tissues where they may re-emerge and spread to susceptible individuals ^2–4^. Although these viruses often exhibit low pathogenicity, their long-term effects, particularly in relation to chronic health conditions, remain a topic of ongoing investigation.

Recent studies have begun to explore the potential links between various viral infections and neurodegenerative diseases (NDDs) including Alzheimer’s disease (AD), amyotrophic lateral sclerosis (ALS), dementia, vascular dementia, and multiple sclerosis MS. Findings suggest that viruses, even those with mild acute pathogenicity, may contribute to alterations in protein levels in neurons and facilitate neurodegeneration over time ^5–7^. Moreover, inflammation caused by viral infections may also contribute to NDDs through direct effects on neuronal function, mitochondrial dysfunction, or via epigenetic changes. Post-mitotic mature human neurons possess robust mechanisms to protect themselves against cell stress and apoptosis ^8^. Therefore, neurons that survive viral infections may persist in a dysfunctional state that influences NDD pathogenesis not only through acute inflammatory damage but also by driving persistent changes in neuronal protein networks and function.

Here we focus on five neurotropic, endemic viruses known for their mild pathogenicity: Herpes simplex virus 1 (HSV-1), Human coronavirus 229E (HCoV-229E), Epstein-Barr virus (EBV), Varicella-Zoster virus (VZV), and Influenza A virus (H1N1). These viruses are widespread and their persistent presence in human populations makes them ideal candidates for studying long-term effects on neuronal health. Given the widespread nature of these viruses, we hypothesize they may influence the protein profiles of neurons, contributing to neurodegenerative processes over time, particularly in older adults. We performed a protein profile analysis on human induced pluripotent stem cell (iPSC)-derived neurons exposed to varying viral loads of these five viruses over periods of one, two, and five days. We identified specific proteins implicated in NDDs that had altered expression levels post-infection. Additionally, all protein levels were cataloged in an accessible database for the scientific community.

Our results demonstrate that exposure to these endemic viruses leads to significant alterations in protein levels in neuronal cells. Pathway analysis revealed that some dysregulated proteins caused by viral infection are associated with neurodegeneration. Although these viruses are typically non-pathogenic, and many individuals experience only mild symptoms after infection, our findings suggest that viral exposures may increase susceptibility to NDDs, particularly in older individuals with multiple viral exposures over their lifetime.

## Results

### Overview of Study Workflow

The human-derived iPSC cell line KOLF2.1J is an ideal model to study neurological disorders. KOLF2.1J iPSCs can be readily differentiated into cells resembling young un-infected neurons, and are free of most genetic variants associated with neurological disorders ^5^. We can study the direct impact of the virus on neurons derived from this iPSC line in a controlled and consistent manner. In this study, KOLF2.1J iPSCs were differentiated into neurons for 21 days and then infected with each of the five common endemic human viruses: HSV-1, HCoV-229E, EBV, VZV, and H1N1, respectively. A single virus can generate hundreds to thousands of copies in an infected cell, enabling the infection of neighboring cells, with viral loads in blood reaching thousands of copies per milliliter. While the blood-brain barrier provides protection for neurons, neurotropic viral infections could expose neurons to hundreds of viruses during the acute phase of infection, depending on the virus type and the severity of infection. To mimic lingering viral loads after acute infection, we selected a range of virus-to-cell ratios (MOI) from low (i.e., 1:10) to high (i.e., 5:1) to infect differentiated neurons. Cells were exposed to viruses for two hours before the media was replaced. Samples were collected one, two, and five days post-infection for proteomic analysis. We conducted a comprehensive large-scale mass spectrometry (MS)-based proteomic analysis of these infected neurons. This approach allowed us to systematically evaluate virus-induced changes in protein expression and identify molecular pathways affected within the infected neuronal cells (Figure 1A). Additionally, we developed a Web app of entire datasets for browsing and visualization (https://niacard.shinyapps.io/CARD_virus/).

**Figure 1.**
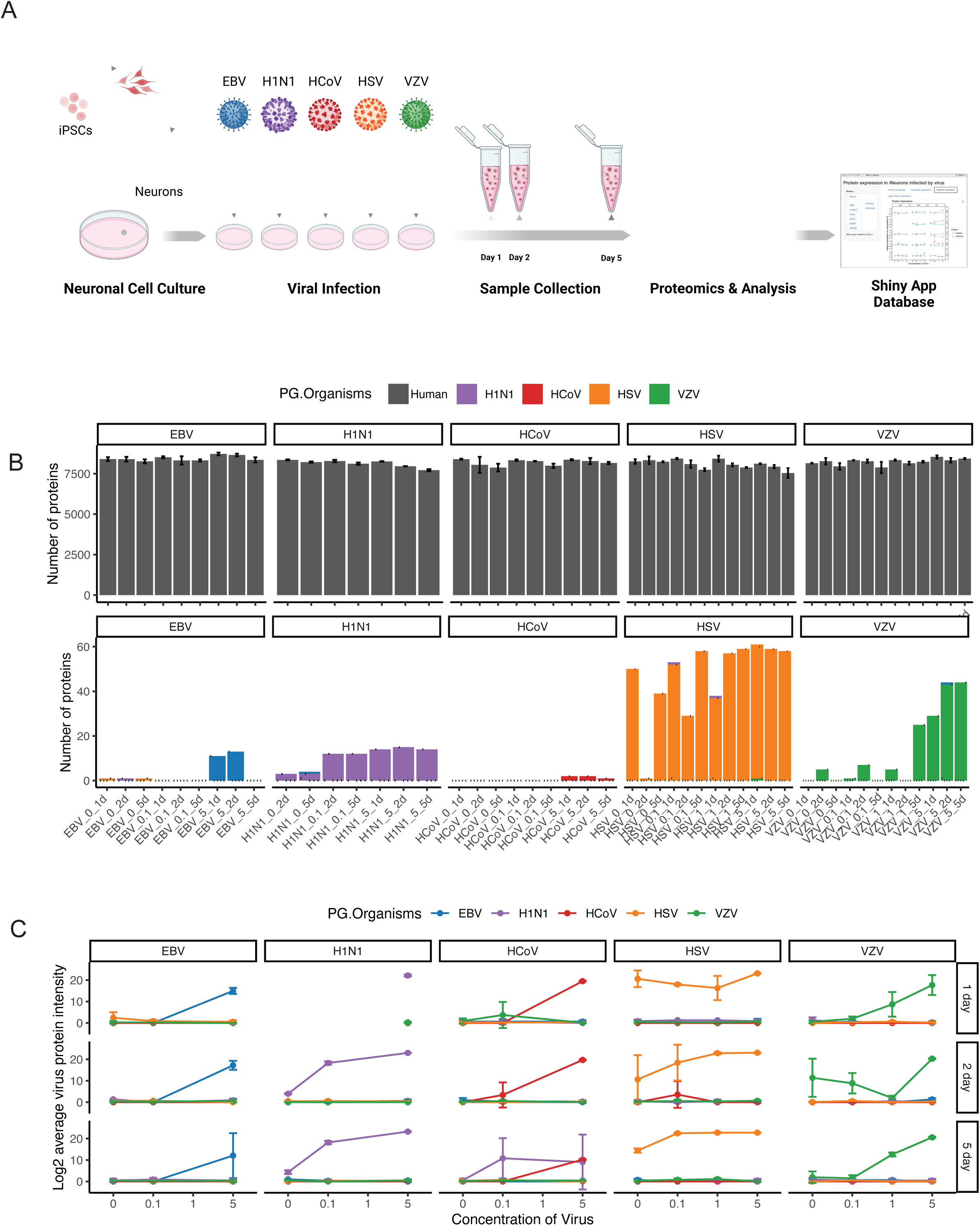
Customized FASTA Database Reveals Virus-Specific Peptides in Infected Neuronal Cultures. (A) Schematic workflow of the study design. Human induced pluripotent stem cells (iPSCs, KOLF2.1J line) were differentiated into. After 26 days of induction, neurons were infected with various human viruses (HSV-1, HCoV-229E, EBV, VZV, and H1N1) at different concentrations. Samples were collected at 1, 2, and 5-days post-infection. Proteomic analyses using mass spectrometry were conducted to identify differentially expressed proteins, assess virus-specific effects, and investigate associated pathways. A Webapp was developed for data browsing (https://niacard.shinyapps.io/CARD_virus/). (B) Human and virus proteins were identified in neuronal cultures using mass spectrometry-based proteomics. (C) Detection and confirmation of virus-specific proteins in infected neuronal samples. Several proteins of interest showed alterations in expression across virus exposure, viral concentration, and exposure time.

### Customized library reveals virus-specific peptides in infected neurons

To enable the detection of viral proteins alongside endogenous cellular proteins, we constructed a customized proteome library containing both viral and human protein sequences. This allowed us to identify approximately 7,500 human proteins (Figure 1B, Figure S1A, Table S1) as well as viral proteins in the infected samples (Figure 1B, Table S2). Notably, we observed an increase in the numbers and intensities of viral proteins and peptides following infection, confirming successful viral infection in the neuronal cultures (Figure 1C). Furthermore, the viral peptides detected were almost entirely unique to each specific virus type, enabling clear differentiation between infections caused by different viruses. These findings also revealed alterations in protein expression across conditions of viral exposure, viral concentration, and exposure time. We conducted cluster analysis to investigate the global impact of viral infections on the proteomic landscape, revealing distinct clustering patterns of proteomic data based on the viral infection conditions (Figure S1B). These results indicate that different viral exposures result in distinct proteomic profiles in neurons.

### Differential Expression Analysis of Common Genes Following Viral Infections

A comprehensive analysis of dysregulated proteins across all virus types and time points revealed key insights into shared and unique responses to viral infections. To identify patterns of protein dysregulation, we performed differential expression analysis of human proteins in neurons infected with the highest viral dosage compared to controls for each virus and time point (Table S3 and S4). We defined dysregulated proteins as those with absolute log2 fold change (FC) greater than 1 and Benjamini-Hochberg adjusted p-value less than 0.01. First, we quantified the number of dysregulated proteins across virus types, revealing the extent of protein alterations caused by each viral infection (Figure 2A). Notably, HSV-1 and H1N1 infections were associated with a higher number of downregulated proteins, whereas HCoV-229E, EBV, and VZV infections induced more upregulation of proteins. Among the viruses, EBV and VZV demonstrated the most extensive protein dysregulation (average of 174 and 186 dysregulated proteins, respectively), while HSV-1, HCoV-229E, and H1N1 exhibited fewer changes (77, 48, and 104 dysregulated proteins, respectively).

**Figure 2.**
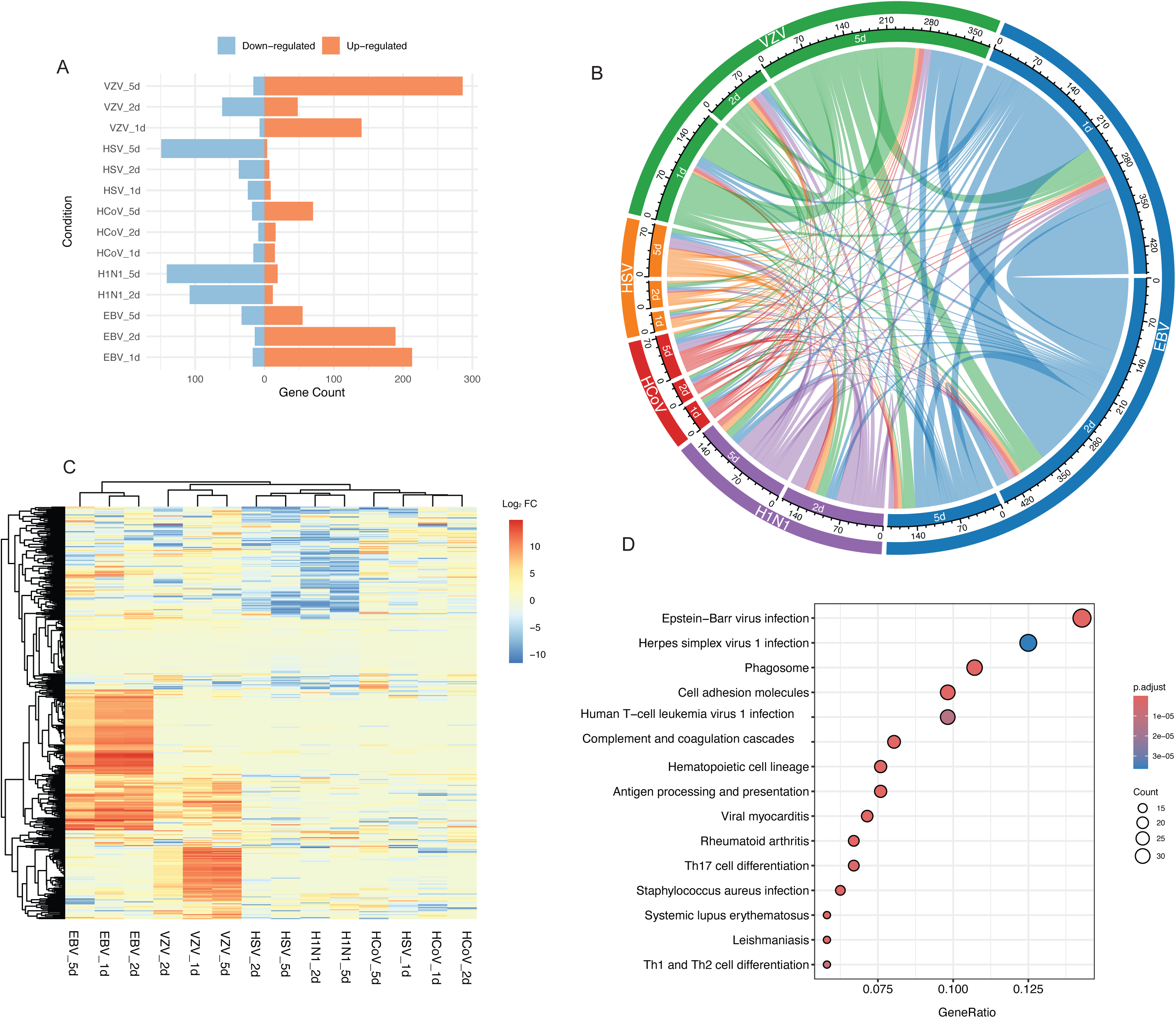
Differential Expression Analysis of Common Genes in Viral Infections. (A) The dysregulated proteins identified in each comparison across different virus types and time points, as identified by gene count, providing an overview of the extent of protein changes. (B) A circular visualization displaying the relationships and overlaps between the dysregulated proteins across different viral infection conditions and time points, as noted by color. (C) A heatmap showing the changes in protein expression of dysregulated proteins across various virus types, highlighting specific patterns of protein alterations in response to each virus. (D) Pathway analysis of the dysregulated proteins shown in (C), revealing the pathways and processes that were significantly altered by viral infections in neuronal cultures.

To further explore these alterations, we assessed the relationships and overlap between dysregulated proteins across the viral conditions. This analysis revealed shared patterns of protein dysregulation. For instance, many dysregulated proteins were consistently shared within the same viral infection across different time points (Figure 2B). Fisher’s exact test confirmed significant overlap in the dysregulated proteins between EBV and VZV-infected neurons (p = 1.26e-22) with an estimated odds ratio of 4.04 (95% CI: 3.11-5.20), suggesting that proteins dysregulated in EBV-infected neurons are about four times more likely to be dysregulated in VZV-infected neurons. This suggests that EBV and VZV neuronal infections may affect common molecular mechanisms or pathways. Similarly, HSV-1 showed notable overlap with H1N1 (p = 9.02e-18, OR: 4.68, 95% CI: 3.38-6.39), indicating potential commonalities in their impact on neuronal proteomes. These findings emphasize the complex interplay of shared and virus-specific proteomic changes in response to different infections (Figure 2C).

To identify the top biological pathways and processes affected by viral infections in neurons, we performed pathway analysis on the significantly altered human proteins shared across different viral infections (Figure 2D, Table S4). Notably, the top two enriched pathways are associated with EBV infection and HSV-1 infection, partially supporting that we are observing virus-specific proteomic changes. Additionally, immune response pathways, including antigen processing and presentation, viral myocarditis were significantly enriched, with key molecules such as HLA, B2M, and TAP1/2, suggesting that viral infections trigger immune-related processes in neuronal cells. These findings provide critical insights into the molecular mechanisms influenced by viral infections in the neuronal environment.

### Dysregulation of Virally-induced Anti-apoptotic and Cognitive Genes in Neurons

Throughout the experiment, we monitored cell morphology daily. No significant changes in morphology, cytopathic effects (CPE), or cell death were observed within one to five days post-infection (data not shown). The neurons maintained their characteristic morphology, with distinct cell bodies and multiple long branches. Therefore, we collected virally-induced dysregulated proteins (i.e., FC >1.5 or FC <0.67) involved in the neuronal anti-apoptotic pathways (Figure 3A). Mature post-mitotic human neurons naturally regulate the expression of anti-apoptotic B-cell lymphoma-2 (BCL-2) family of proteins to prevent cell death and withstand cellular stress. Notably, we observed that BCL-2 levels were significantly increased in neurons infected with EBV and HCoV-229E. VZV also upregulated anti-apoptotic proteins, including ANGPT1, BDNF, CCL2, IL6ST, NRP1, and SEMA3E. In contrast, HSV-1 infection did not significantly increase anti-apoptotic protein levels. We hypothesize that the intricate interplay of these proteins may contribute to neuronal survival during infection. Furthermore, we visualized dysregulated proteins involved in cognition and learning or memory pathways (Figure 3B and 3C). Our results suggested that endemic viruses may differentially impact cognitive function and learning ability.

**Figure 3.**
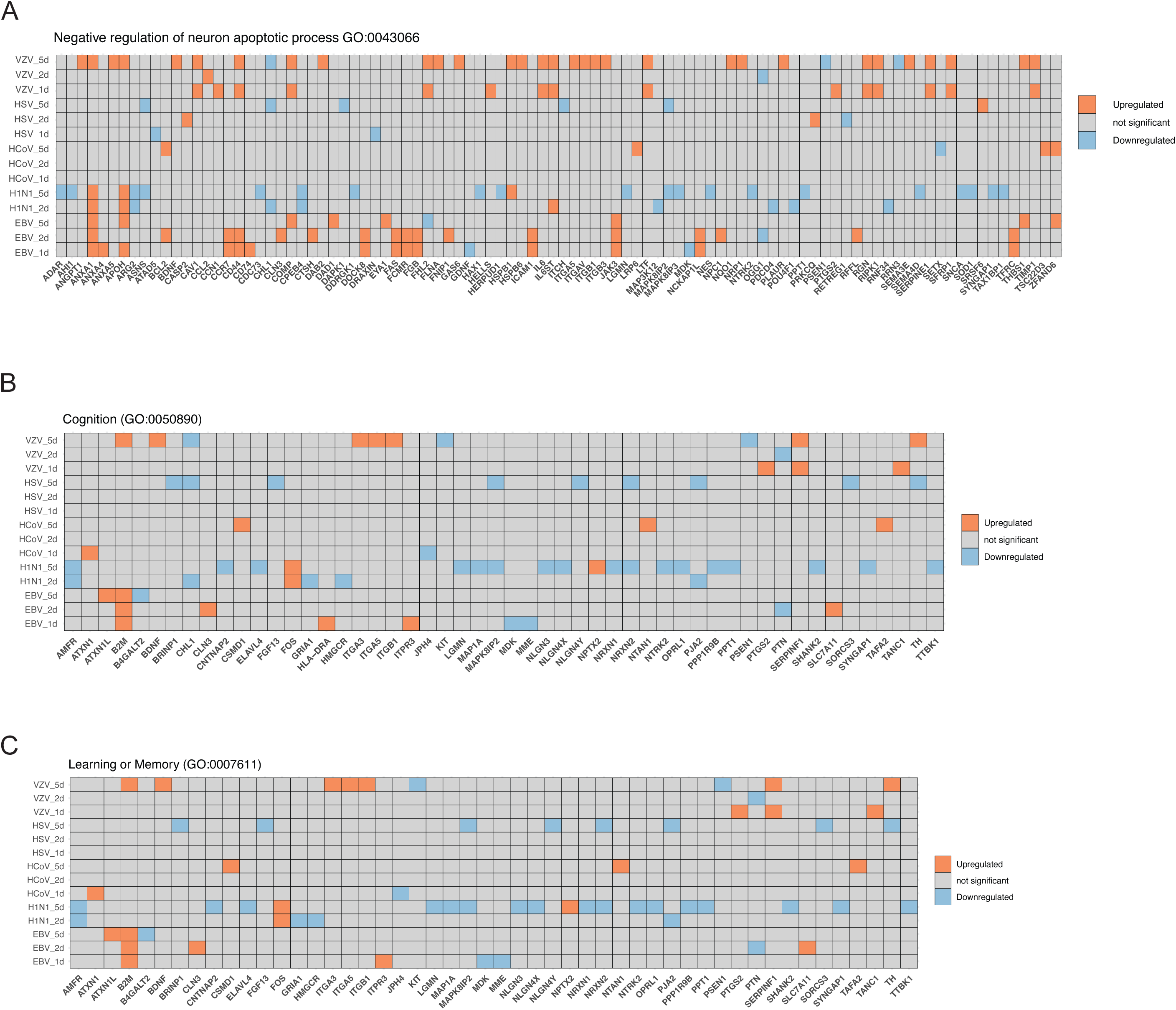
Pathways associated with neuron apoptosis and cognition. (A) The significant dysregulated proteins enriched in the negative regulation of neuron-apoptotic process (GO:0043524). Proteins were considered upregulated with log2 FC > 0.585 (adj. p-value < 0.05) and downregulated with log2 FC < −0.585 (adj. p-value < 0.05). (B) The significant dysregulated proteins enriched in the Cognition (GO:0050890). Proteins were considered upregulated with log2 FC > 0.585 (adj. p-value < 0.05) and downregulated with log2 FC < −0.585 (adj. p-value < 0.05). (C) The significant dysregulated proteins enriched in the anti neuron-apoptotic process (GO:0043524). Proteins were considered upregulated with log2 FC > 0.585 (adj. p-value < 0.05) and downregulated with log2 FC < −0.585 (adj. p-value < 0.05).

### WGCNA Reveals Virus-Specific Gene Modules and Their Functional Associations

To investigate the common molecular mechanisms underlying virus-induced neuronal dysfunction, we performed Weighted Gene Co-expression Network Analysis (WGCNA) using all human proteins across all conditions. This analysis generated clusters of co-expressed genes, referred to as gene modules, uniquely associated with viral infection, revealing cellular processes and molecular pathways altered in response to viral exposure. Using WGCNA, we identified 18 distinct modules of human proteins highly correlated with one another across all the conditions (Figure 4A, Figure S2A, Table S5). Among these, module 10 (1407 proteins), module 11 (854 proteins), and module 5 (578 proteins) contained the largest number of proteins (Figure S2B). When we correlated the relationship between these modules and viral protein abundance in neurons using a module-trait association analysis, we found that many modules are significantly associated with the expression of viral proteins (Figure 4B). Seven modules displayed correlations with absolute r values ranging from 0.4 to 0.67 for at least one virus, all with pLJ<LJ0.001, calculated using Pearson’s correlation with nLJ=LJ145). VZV proteins were positively correlated and HSV-1 proteins were negatively correlated with the majority of the modules, highlighting their potential involvement in virus-induced changes in neurons (Figure 4B).

**Figure 4.**
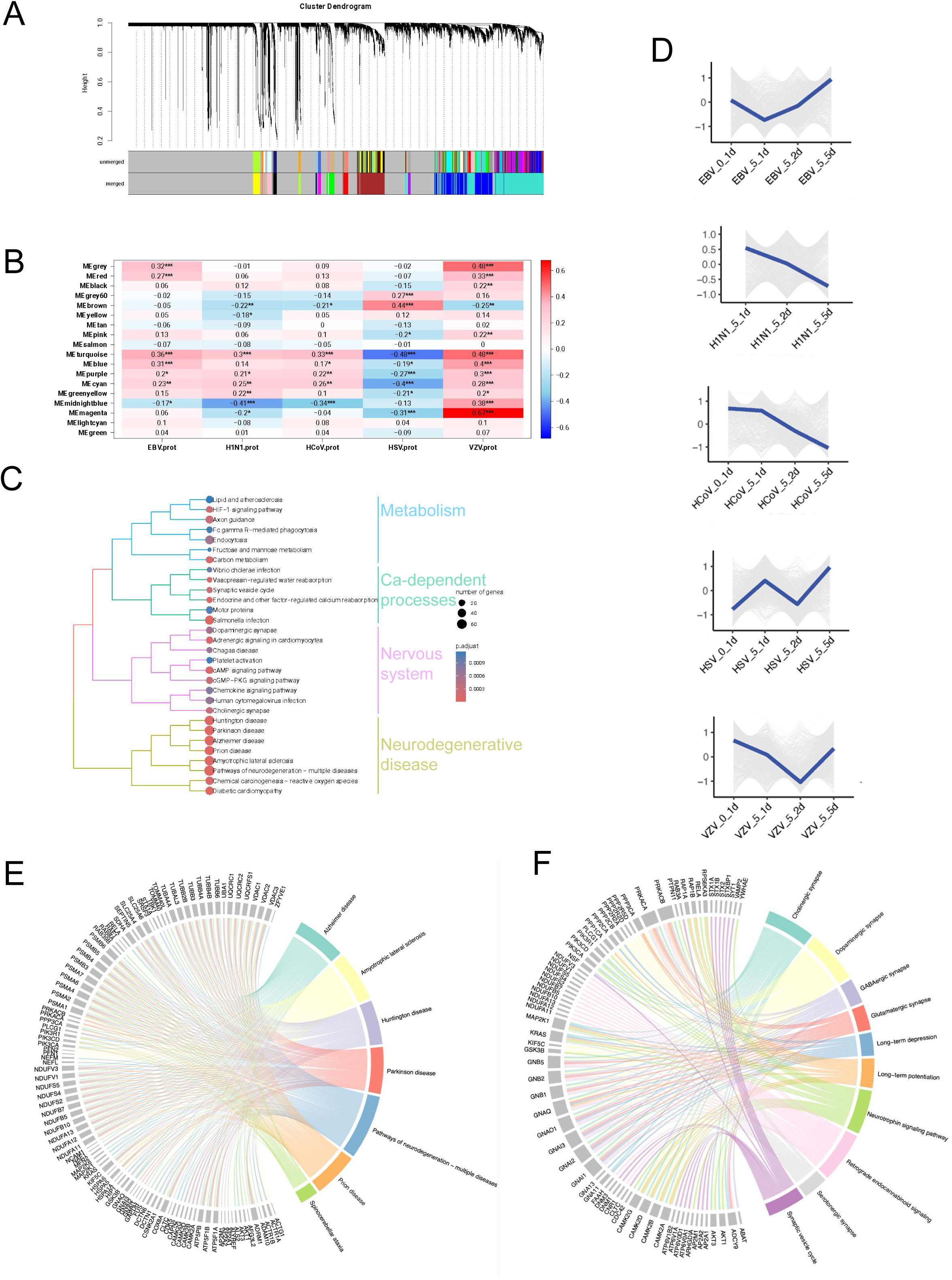
WGCNA of Virus-Related Gene Modules. (A) Gene dendrogram obtained through average linkage hierarchical clustering, with the color row underneath the dendrogram showing the module assignment determined by the Dynamic Tree Cut method. The proteins are clustered in 18 modules based on protein abundance in virus infected neurons. (B) Module-trait association analysis, where each row corresponds to a gene module and each column corresponds to a trait. Each cell contains the corresponding p-value from the linear mixed-effects model, indicating the relationship between modules and traits. *p<0.05, **p<0.01, ***p<0.001, ****p<0.0001 (C) KEGG pathway analysis of the module 11 proteins, highlighting its association with nervous system function and NDDs. (D) Chord diagram depicting the module 11 proteins involved in seven neurological disorders identified through enrichment analysis. (E) Proteins expression of the module 11 in neurons across different time points and virus types post-infection. (F) Chord diagram depicting the module 11 proteins involved in ten nervous system pathways, as identified in the enrichment analysis.

To investigate the functional significance of the identified modules, we performed pathway analysis for each module. Among all modules, module 11 contained 854 co-expressed human proteins and emerged as particularly noteworthy due to the large number of proteins within the module and the strong association with viral protein expression (Figure S2B). Pathway analysis of this module revealed that eight pathways are directly related to NDDs (Figure 4C). We identified proteins strongly linked to multiple NDDs, including 60 proteins associated with Parkinson’s disease, 63 with AD, 57 with prion disease, 64 with ALS, and 57 with Huntington’s disease (Figure 4D). Moreover, the five viruses we tested caused unique changes in the trajectories of module 11 proteins, suggesting that viral infections may contribute to neurodegenerative processes in different ways (Figure 4E, Figure S2C). A protein–protein interaction network of NDD-related genes highlighted interactions among proteins dysregulated during viral infections (Figure S2D).

Additionally, proteins of module 11 were significantly enriched in pathways essential to nervous system function, such as cholinergic synapses and synaptic vesicle cycling (Figure 4C, Figure 4F). These findings highlight the potential of viral infections to influence key molecular pathways underlying neuronal health and neurodegeneration. In addition to module 11, other modules also significantly correlated with viral infection demonstrated distinct biological significance, shedding light on various cellular processes impacted by viral infections. The module 16 was significantly enriched in pathways related to wound healing, emphasizing its role in tissue repair and regeneration (Figure S3A). This aligns with evidence that viral infections can modulate immune responses and cellular signaling to influence tissue repair processes, either promoting recovery or exacerbating damage ^9,10^. The module 10 was associated with protein degradation pathways, including lysosome and ubiquitin-proteasome systems, which are crucial for cellular protein turnover (Figure S3B). Viruses are known to exploit these pathways to degrade host proteins or evade immune surveillance, often resulting in broader disruptions to cellular homeostasis ^11,12^. The module 5 was enriched in pathways involved in DNA repair and cell cycle regulation, emphasizing its role in maintaining genomic stability and cellular integrity (Figure S3C). DNA viruses are known to interfere with these processes, creating a replication-favorable environment while potentially inducing genomic instability ^13,14^. The module 2 showed enrichment in pathways related to mRNA splicing and processing, focusing on RNA metabolism and regulation (Figure S3D), which viruses often manipulate to control host gene expression and optimize conditions for their replication ^15,16^. These findings collectively highlight the intricate involvement of virus-co-expressed proteins in critical cellular pathways, offering insights into how viral infections disrupt essential biological processes and contribute to cellular dysfunction, repair mechanisms, and broader disease pathology.

### Validation of Virus-Dysregulated Proteins in Alzheimer’s Disease Patient Samples Using Proteomics Data

We further focused our study on AD due to its potential association with viral infection. To gain a deeper understanding of virus-dysregulated proteins and their potential links to AD, we examined whether these proteins are present in Alzheimer’s disease patient samples. We compared differentially expressed proteins (DEPs) caused by viral infection with in vivo observations from seven publicly available studies. These studies included MS-based proteomics data from postmortem brain tissues and cerebrospinal fluid (CSF) of AD patients and healthy controls ^17^.

We found 22 overlapping DEPs associated with viral infection in neurons were significantly changed proteins in the AD human studies (Figure 5A). The higher abundant proteins in the AD brain were also observed in upregulated proteins associated with VZV infection (i.e., 9 and 11 DEPs on day1 and day5, respectively). Similarly, 5 of DEPs associated with EBV infection were also observed with higher abundance in the AD brain. While the AD associated proteins observed in the CSF are not correlated with those in the brain. These results indicate a strong association of viral infection with AD pathogenesis *in vivo*. Furthermore, a PPI network highlighted shared dysregulated proteins during viral infection and AD, highlighting potential molecular intersections between viral exposure and AD pathogenesis (Figure 5B). AD-related pathways involved in ICAM1 ^18^, MYLK ^19^, C4A ^20^, SCARB1 ^21^, HSPB1 ^22^,CD44 ^23^, IGFBP5 ^24^, PTX3 ^25^,CPT1A ^26^, EFR3A ^27^,DECR1 ^28^,CEMIP ^29^, and SMOC1 ^30^, are also dysregulated by viral infections, suggesting a direct impact of viral exposure on key pathways involved in AD. These findings provide novel insights into the potential role of viral infections in influencing pathways relevant to AD and neurodegeneration, underscoring the need for further investigation into how viruses may exacerbate or initiate AD pathology.

**Figure 5.**
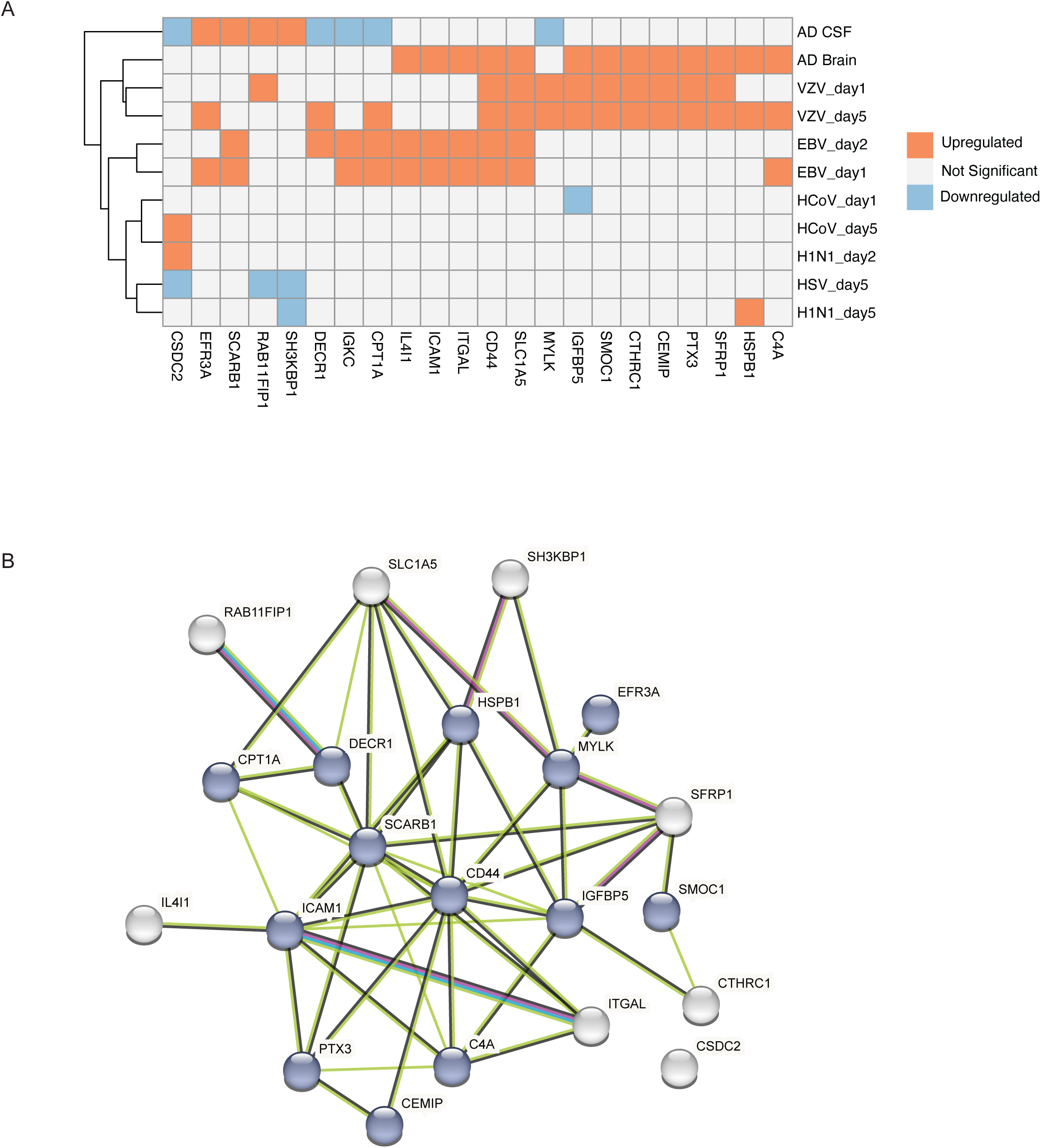
Correlation of Virus-Induced Protein Changes with Alzheimer’s Disease. (A) Heatmap displaying the correlation of altered proteins by viral exposure in neurons and their abundance alternation in the cerebrospinal fluid (CSF) and postmodern brain from AD patients compared to healthy controls. Proteins were considered upregulated with log2 FC > 0.585 (adj. p-value < 0.05) and downregulated with log2 FC < −0.585 (adj. p-value < 0.05). (B) Protein–protein interaction (PPI) network of shared dysregulation of proteins during viral infection and AD.

## Discussion

Our study provides critical insights into how common viral infections impact neuronal function. We used iPSC-derived neurons and a proteomic approach to identify virally-induced changes in protein expression. We observed distinct, virus-specific proteomic alterations, with some viruses — such as EBV and VZV — causing extensive dysregulation while others, like HSV-1 and H1N1, induced more subtle shifts. These findings highlight the diverse ways in which viruses interact with host neurons, potentially contributing to neurodegenerative disease risk.

By examining five endemic neurotropic viruses — HSV-1, HCoV-229E, EBV, VZV, and H1N1 — we aimed to determine whether mild, persistent infections could subtly alter neuronal protein networks in ways that predispose cells to long-term dysfunction. Given that these viruses collectively infect billions of people worldwide, their potential role as hidden contributors to neurodegeneration warranted closer examination. Indeed, we found that viral infection rapidly reprogrammed neuronal proteomes within just one to five days post-infection. To understand the function of the proteins co-expressed with viral proteins, we performed further pathway analysis. This analysis revealed that some of the dysregulated proteins are associated with pathways related to the nervous system, including those implicated in Alzheimer’s, Parkinson’s, and Huntington’s diseases. Despite neuronal survival, these molecular alterations suggest a slow-burn effect where transient infections may prime neurons for dysfunction long before overt degeneration occurs. The survival of most cells suggests resistance to acute cytotoxicity from these viruses, potentially due to intrinsic survival mechanisms, the latency-prone nature of certain viruses, or the absence of immune-mediated damage. Additionally, we used low viral loads to mimic persistent infections rather than acute-phase viral replication. Certain viruses modulate apoptosis and autophagy to prolong host cell survival and facilitate persistent infections. Viral infections have been shown to impact neuronal function, with recent studies demonstrating the ability of viruses to infect neurons and alter cellular responses. For example, a recent study on SARS-CoV-2 infection in neurons revealed significant transcriptional and proteomic changes, indicating that viral infections can influence neuronal homeostasis ^31^. A study by Readhead et al. demonstrated that HSV-1 infection in trigeminal ganglion neurons leads to widespread transcriptional and epigenetic changes ^32^. However, these studies primarily focused on COVID-19 and HSV-1 without linking infection-induced molecular signatures to specific neurodegenerative diseases. In contrast, our findings suggest that viral infections may drive proteomic alterations in neurons that resemble those observed in AD patient samples. This highlights the potential for viral infections to contribute to neurodegenerative disease mechanisms, warranting further investigation into the long-term consequences of virus-induced neuronal dysregulation.

To the best of our knowledge, this is the first study to have identified virally-induced dysregulated proteins linked to neurodegenerative pathways in an iPSC-derived neuronal model. The WGCNA revealed that proteins in the module 11, in particular, were associated with NDDs such as AD, Parkinson’s, and Huntington’s disease. Pathway analysis further underscored that viral infections may exacerbate these conditions by modulating key processes related to neuronal health, such as synaptic function, protein degradation, and immune response. The overlap of virus-dysregulated proteins with those implicated in AD, as demonstrated through our correlation analysis, suggests a potential link between viral infections and the onset or progression of neurodegeneration. These findings underscore the need to further investigate how viral persistence, immune responses, and neuronal stress pathways interact to drive infection-related neurodegeneration. Future studies should explore whether these virally-induced proteomic changes have lasting consequences for neuronal function and survival and assess the role of neuroimmune interactions in amplifying these effects. This study provides a compelling argument for further exploring the role of viral infections in modulating neurodegenerative disease pathways, particularly in individuals with a history of viral exposure.

Additionally, the results emphasize the importance of considering the broader cellular context in understanding viral infections in the nervous system. Beyond the module 11, other co-expression modules related to DNA repair, tissue repair, and protein degradation were also significantly enriched in pathways associated with viral infection. These findings suggest that viral infections not only trigger immune responses but also interfere with critical cellular processes such as protein turnover and DNA repair. These disruptions may contribute to the long-term dysfunction of neurons, further highlighting the potential risks of viral exposure in the context of NDDs. Our results serve as an important bridge between epidemiological associations and molecular mechanisms, providing cellular-level evidence that viral infections may act as early initiators of neurodegenerative pathways rather than merely exacerbating preexisting conditions. However, the significant proteomic disruptions we observed indicate that infection may subtly compromise neuronal homeostasis, increasing susceptibility to later insults or accelerating disease processes over time.

In conclusion, our study reveals the intricate ways in which viral infections influence neuronal proteomes, offering new insights into the molecular mechanisms underlying virus-induced neurodegeneration. By linking viral infection to well-established pathways in NDDs, these findings open up new avenues for research into the role of viral infections in the onset and progression of diseases like AD. Further studies are needed to explore how these virally-induced changes in protein expression might be targeted therapeutically to prevent or mitigate the impact of viral infections on neuronal health.

## Acknowledgement

This research was supported in part by the Intramural Research Program of the NIH, National Institute on Aging (NIA), National Institutes of Health, Department of Health and Human Services; project number ZIAAG000534. This work was also supported by the Intramural Research Program at the National Institute of Environmental Health Sciences (ZIC ES102506-09). We thank the NIH HPC system (*Biowulf*, http://hpc.nih.gov) for making this work possible. We also thank Drs. Kaoru Inoue and Carl Bortner for critical reading of the manuscript.

## Data availability

The MS raw files of this study are deposited to the ProteomeXchange Consortium via the PRIDE ^33^ partner repository with the dataset identifier PXD061286. The MS raw files are referred to in original publications. The processed protein abundance data can be found in the supplementary data.

## Author contributions

Conceptualization, N.P.M., S-H.C., Z.L., J.L.Y., Y.A.Q.

Methodology, Z.L., D.C., Y.H., J.E., N.P.M., S-H.C., Y.A.Q., D.M.R.

Data curation, Z.L., J.E., Y.H., C.W., M.A.N., Y.A.Q.

Formal analysis, Z.L. D.M.R., C.W., Y.H., D.M.R., J.E., Y.A.Q.

Investigation, Z.L., N.P.M., L.F., M.R.C., A.B.S., Y.A.Q.

Resources, N.P.M., J.L.Y., C.P., K.M.A.

writing – original draft, Z.L., N.P.M., S-H.C, Y.A.Q.

writing – review & editing, all authors.

visualization, Z.L., J.E., P.J., M.A.N., C.W.

supervision, N.P.M., L.F., M.A.N., M.R.C., A.B.S., Y.A.Q, J.L.Y.

## STAR Methods

### Key resources table

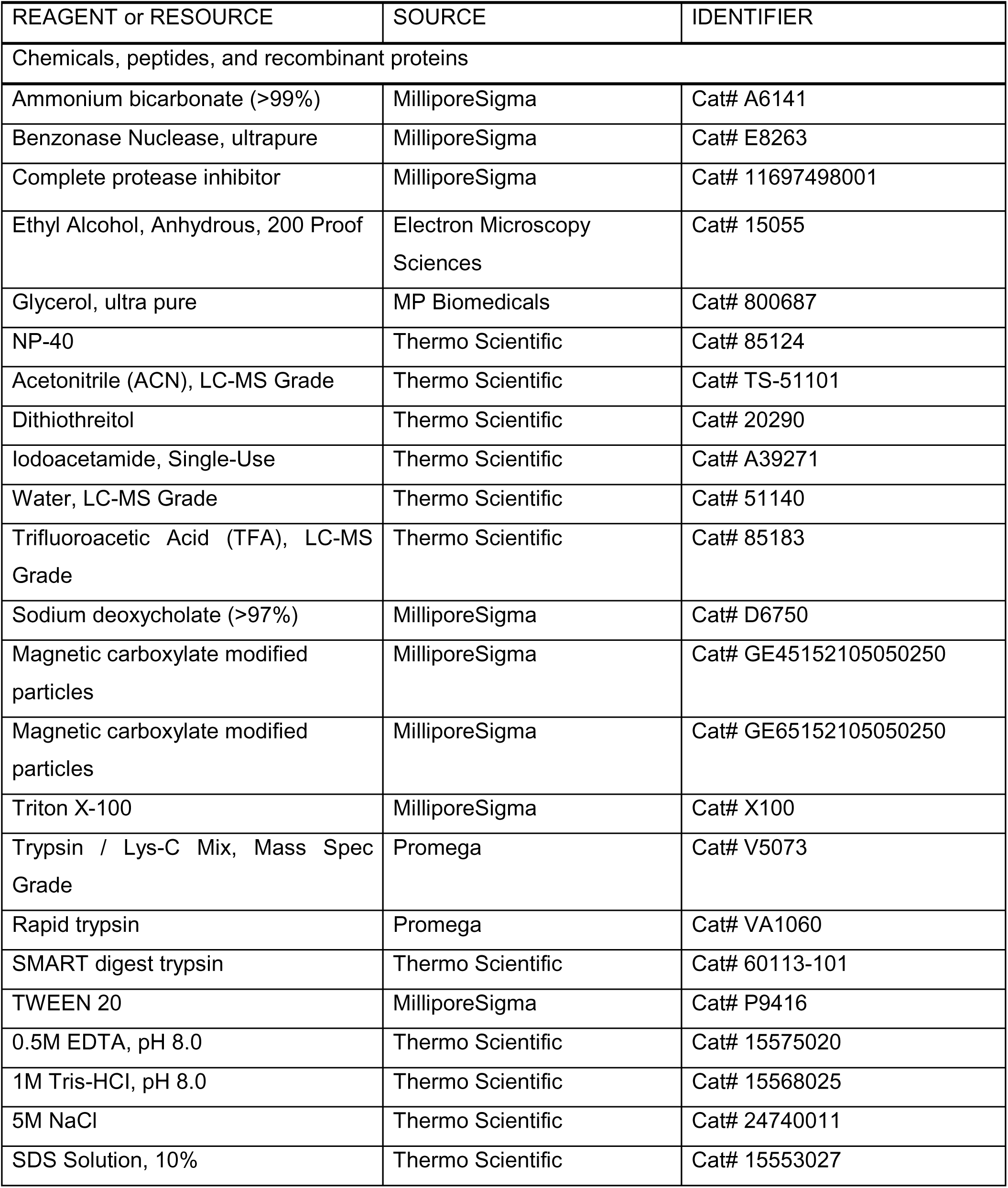
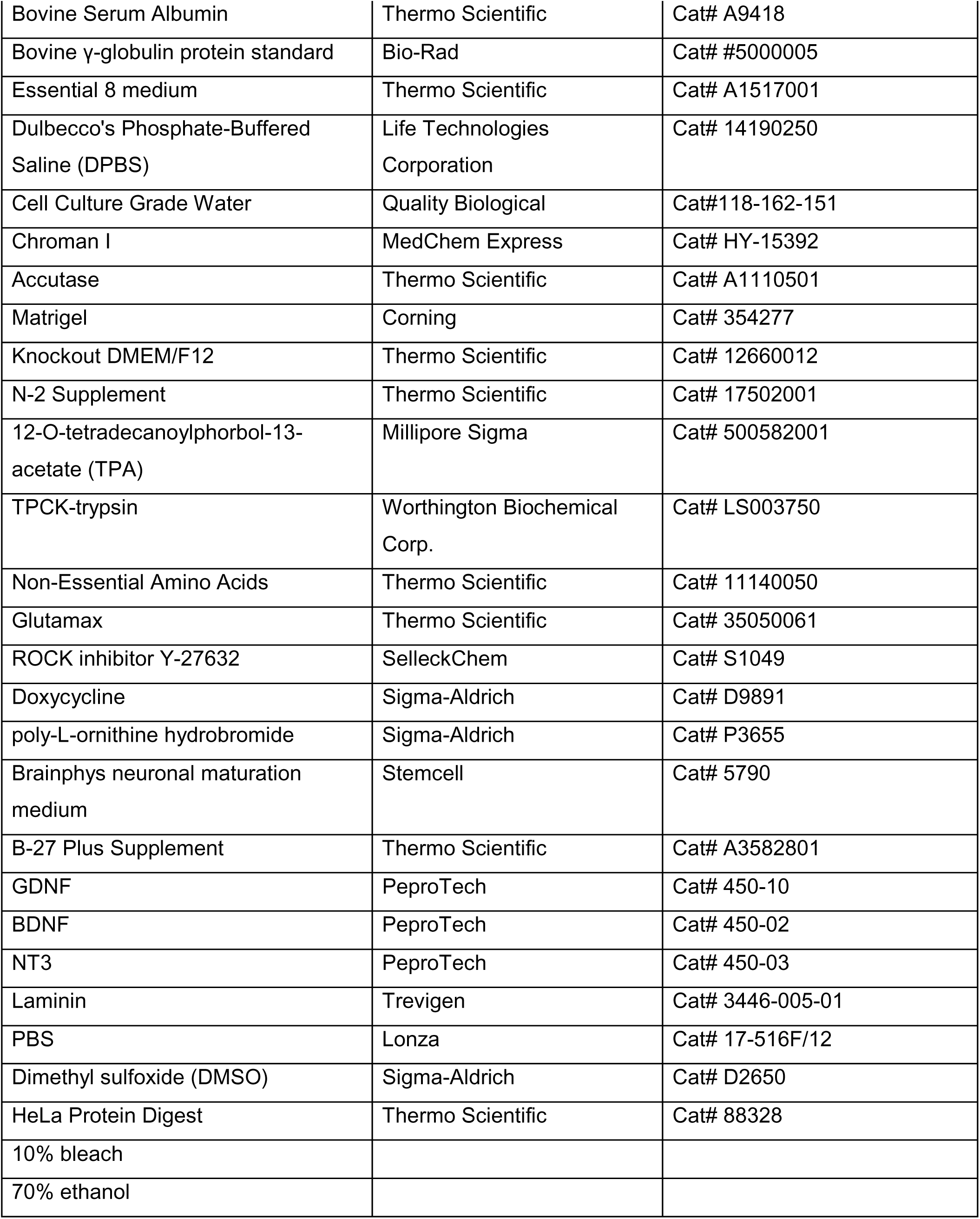
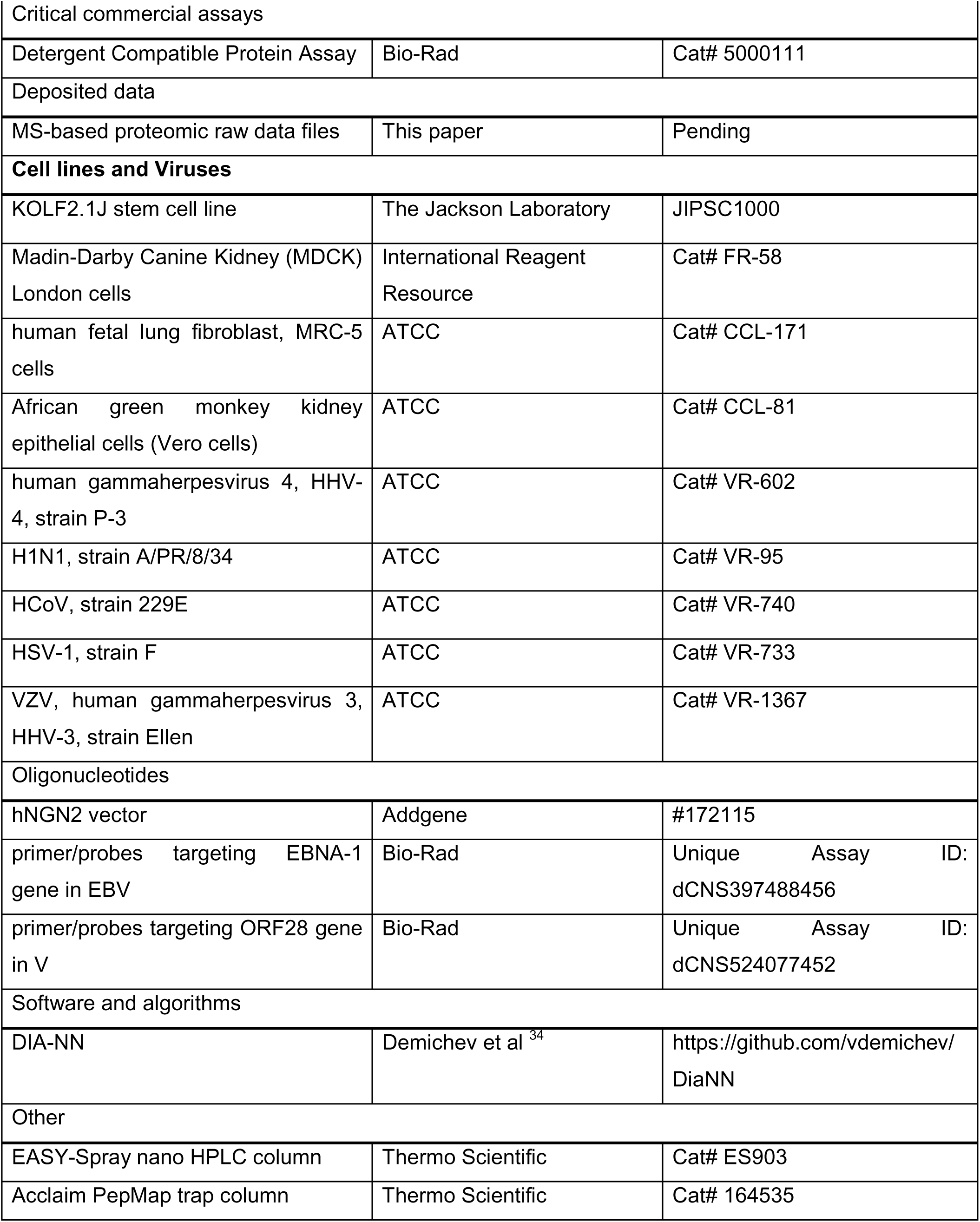
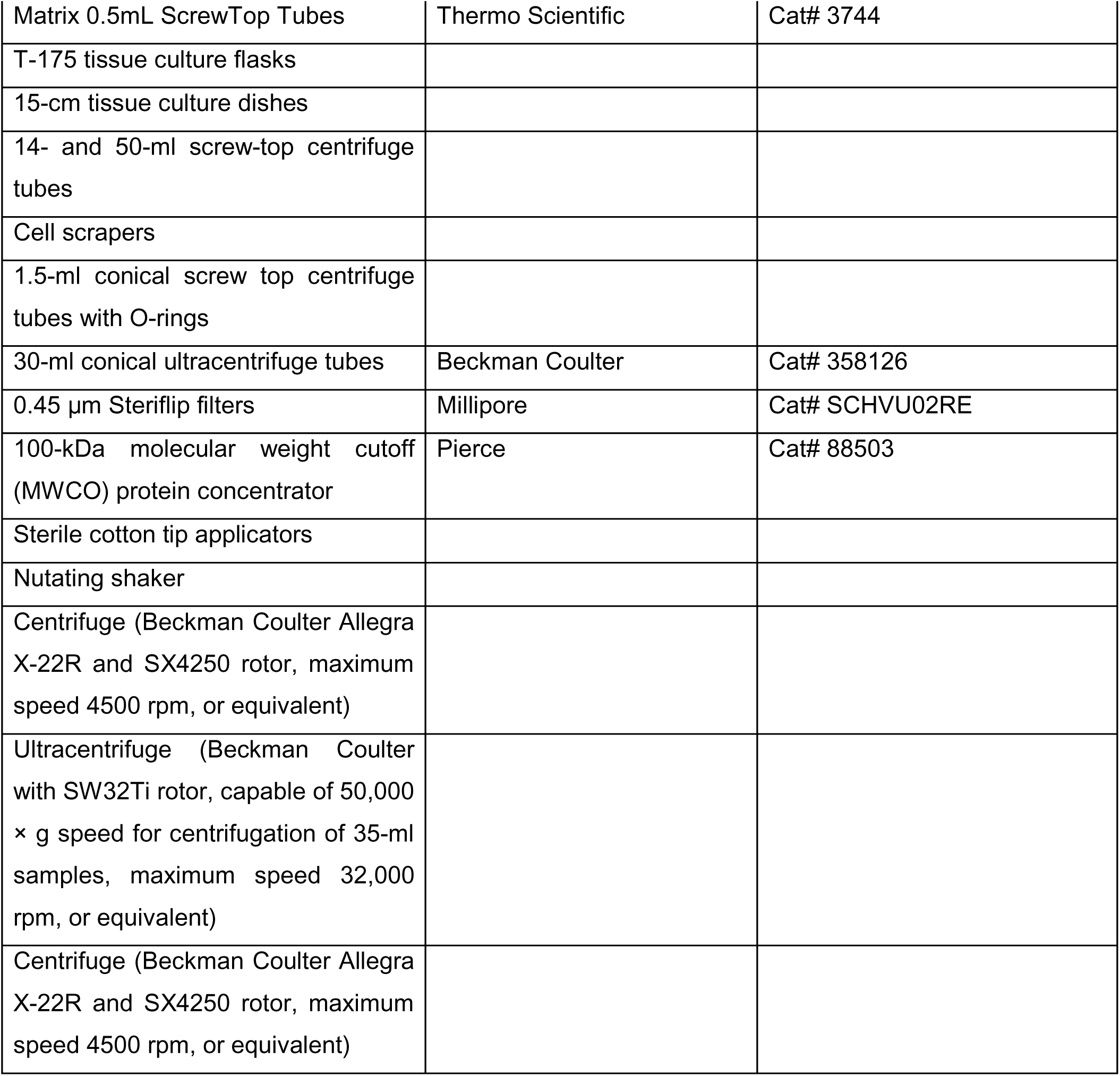

## Resource availability

### Lead contact

Further information and requests for resources and reagents should be directed to and will be fulfilled by the lead contact, Yue Andy Qi.

### Materials availability

This study did not generate new unique reagents.

### Experimental model and subject details

In this study, we used the KOLF2.1J line derived from a caucasian male, obtained from a reference parental line of the iNDI project (The Jackson Laboratory, https://www.jax.org/jax-mice-and-services/ipsc). Material transfer agreement between the Jackson Laboratory and the Center for Alzheimer’s and Related Dementias at the NIH was obtained prior to conducting any experiment. Our previous study reports genetic and proteomic characterization of the KOLF2.1J iPSC line ^35,36^.

## Method details

### Human iPSC Cell Culture and Differentiation into Neurons

The KOLF2.1J cell line, which expresses the human neurogenin 2 (NGN2) transcription factor under a tetracycline-inducible promoter (iNGN), was obtained from the Jackson Laboratory (Cat# JIPSC1000) and cultured as previously described ^35,37^. Briefly, cells were expanded and then transferred to 6-well plates, with each well seeded with 8e5 cells in 2 mL of induction media consisting of Knockout DMEM/F12, 1X N-2 Supplement, 1X Non-Essential Amino Acid, and 1X Glutamax, with 10 µM of ROCK inhibitor Y-27632 and 2 µg/mL Doxycycline. Cells were maintained in induction media until day 4, after which the media was switched to neuronal maturation media consisting of Brainphys neuronal maturation medium, 1X B-27 Plus Supplement, 10 ng/mL GDNF, 10 ng/mL BDNF, 10ng/mL NT3, 1µg/mL laminin and 2 µg/mL Doxycycline. Cells were monitored daily, and on alternate days, half of the media was replaced with fresh, pre-warmed media at 37° C. The cell morphology and survival were monitored under a microscope daily for the entire length of the experiment.

### Viral Transduction

iNGN cells were allowed to differentiate for 21 days to form mature neurons in 6-well plates. Prior to infection, half of the media from each well was removed and incubated separately in sterile 6-well plates at 37°C (referred to as stored Media-A). Cells were infected with viruses at multiplicity of infection (MOI) of 0, 0.1, 1, or 5. All infections were performed using equivalent volumes. Virus diluents were used to normalize the volumes. Cells were incubated with viruses for two hours at 37°C. Then, the entire media in each well was replaced with the aforementioned stored Media-A and supplemented with 1 mL of fresh neuronal maturation media. Cells were harvested at one, two, and five days post-infection.

### Virus Propagation and Treatment

#### Epstein-Barr Virus (EBV) Propagation and Treatment

Human Burkitt’s lymphoma cells infected with Epstein-Barr virus (EBV, human gammaherpesvirus 4, HHV-4, strain P-3) were obtained from ATCC (catalog# VR-602). The cells were cultured in RPMI-1640 medium (ATCC, catalog# 30-2001) supplemented with 15% fetal bovine serum (FBS; Avantor Seredigm).

EBV was harvested and concentrated according to a published protocol ^38^. Briefly, EBV replication was induced using phorbol ester (12-O-tetradecanoylphorbol-13-acetate, TPA, Millipore Sigma catalog# 500582001) at a concentration of 20 ng/mL for 5 days. The cells and media were then harvested and sonicated to release the EBV from within the cells. Cell debris was removed by centrifugation at 1,000 × g for 10 minutes. The supernatant was filtered using 0.45-µm low protein-binding filters.

EBV in the supernatant was concentrated by ultracentrifugation for 2 hours at 75,000 × g, 4°C, using a swinging-bucket rotor. The resulting pellet was resuspended in PBS and used for infections as described in the “Viral Transduction” section of the Methods. Media containing TPA was processed in the same way as the media with virus: it was concentrated, and PBS was added to the tubes for use as a diluent in subsequent experiments. Viral titer was determined using digital droplet PCR (ddPCR) with primer/probes targeting EBNA-1 gene (Bio-Rad, Unique Assay ID: dCNS397488456).

#### Influenza A Virus (H1N1) Propagation and Treatment

H1N1, strain A/PR/8/34, was obtained from ATCC (catalog# VR-95). The virus was propagated in Madin-Darby Canine Kidney (MDCK) London cells (ATCC, catalog# FR-58). MDCK London cells were cultured in high-glucose Dulbecco’s Modified Eagle Medium (DMEM; Gibco) supplemented with 10% fetal bovine serum (FBS; Avantor Seredigm), 4 mM L-glutamine (Gibco), and 1 mM sodium pyruvate (Sigma-Aldrich).

For virus propagation, 1.5e7 MDCK London cells were seeded in T175 flasks and incubated for 24 hours at 37°C. Cells were then infected with influenza A virus at a multiplicity of infection (MOI) of 0.01. Briefly, 5 mL of DMEM containing 1 μg/mL TPCK-trypsin (Worthington Biochemical Corp., catalog# LS003750) was added to the flask containing the virus inoculum and incubated for 1 hour at 37°C to allow viral entry. Following this incubation, an additional 20 mL of DMEM containing 1 μg/mL TPCK-trypsin was added. At 72 to 96 hours post-infection, the supernatant was collected and centrifuged at 2,000 x g for 10 minutes at 4°C. The supernatant was then aliquoted, stored at −80°C, and used for infection as described in the “Viral Transduction” section of the Methods. Viral titers were determined by plaque assay on MDCK London cells. Prior to viral infection, 1 mL of medium was removed from each well. The virus was added to the wells at an MOI of 0.1 or 5, and the plates were incubated at 37°C for 2 hours. After incubation, 1 mL of fresh medium containing 1 μg/mL TPCK-trypsin was added to each well. Cells were collected at the desired time points for further analysis.

#### Human Coronavirus 229E (HCoV-229E) Propagation and Treatment

HCoV, strain 229E, was obtained from ATCC (catalog #VR-740) and propagated in MRC-5 cells (ATCC, catalog #CCL-171), a human fibroblast cell line. MRC-5 cells were cultured in Eagle’s Minimum Essential Medium (EMEM) supplemented with 10% fetal bovine serum (FBS; Avantor Seradigm).

For viral propagation, 1.5 × 10LJ MRC-5 cells were seeded into T175 flasks and subsequently infected with HCoV-229E. After the onset of significant cytopathic effect (CPE), both the supernatant and cells were collected. The mixture was vortexed and centrifuged at 2,000 × g for 10 minutes at 4°C to remove cell debris. To concentrate the virus, the supernatant was ultracentrifuged over a 20% sucrose cushion at 50,000 × g for 2 hours at 4°C. The viral pellet was resuspended in phosphate-buffered saline (PBS). Viral aliquots were stored at −80°C until use, as described in the “Viral Transduction” section. Viral titers were determined using a plaque assay on MRC-5 cells.

#### Herpes simplex virus type 1 (HSV-1) Propagation and Treatment

HSV-1, strain F, was obtained from ATCC (catalog# VR-733). The virus was propagated in African green monkey kidney epithelial cells (Vero cells, ATCC, catalog# CCL-81). Vero cells were cultured in high-glucose Dulbecco’s Modified Eagle Medium (DMEM, Gibco) supplemented with 10% fetal bovine serum (FBS, Avantor Seredigm).

For virus propagation, 1.5e7 Vero cells were seeded in T175 flasks and incubated for 24 hours at 37°C. Cells were then infected with HSV-1 at a multiplicity of infection (MOI) of 0.01. Briefly, 5 mL of DMEM was added to the flask containing the virus inoculum and incubated for 1 hour at 37°C to allow viral entry. Following this incubation, an additional 20 mL of DMEM was added to the flask. At 24 to 48 hours post-infection, when more than 90% of cells displayed cytopathic effects, the cell-virus supernatant was collected and centrifuged at 433 x g for 5 minutes at 4°C. 10 mL of supernatant was removed and retained, leaving the cell pellet in the tube. To release cell-associated viruses, the cell pellet was subjected to sonication in an ultrasound bath (Diagenode Bioruptor, model: UCD-200) until the solution appeared clear. The sonicated solution was then centrifuged at 1,743 x g for 10 minutes at 4°C. The supernatant was transferred to a new tube, aliquoted, and stored at −80°C. Viral titers were determined by plaque assay on Vero cells.

#### Varicella Zoster Virus (VZV) Propagation and Treatment

VZV, human gammaherpesvirus 3, HHV-3, strain Ellen, was obtained from ATCC (catalog# VR-1367) and propagated in MRC-5 cells (ATCC, catalog# CCL-171), a human fetal lung fibroblast cell line. MRC-5 cells were cultured in Eagle’s Minimum Essential Medium (EMEM) supplemented with 10% fetal bovine serum (FBS, Avantor Seradigm).

VZV propagation and concentration followed a previously published protocol ^39^. Briefly, 2.5e6 MRC-5 cells were infected with 0.4 mL of VZV (6.23e4 transducing units/mL) in media containing 2% FBS for 16–18 hours. The following day, the media was replaced with fresh EMEM supplemented with 10% FBS, and the cells were incubated at 37°C for an additional 7 days. Upon reaching a significant CPE, cells were scraped and transferred to media, then used to infect fresh, uninfected MRC-5 cells at a ratio of 1 infected cell to 5 uninfected cells to propagate the stock.

After two weeks, when CPE became prominent, both the cell supernatant and cell pellets were collected. To release cell-associated virus, the cell pellet was subjected to sonication in an ultrasound bath (Diagenode Bioruptor, model UCD-200) until the solution appeared clear. The sonicated sample was then centrifuged at 1,500 × g for 15 minutes at 4°C to remove cellular debris. The supernatant was transferred to a new tube and concentrated using Pierce Protein Concentrators (PES, 100K MWCO, catalog# 88537).

Media without virus was processed in the same manner as the virus-containing media and served as a diluent in subsequent experiments. The resulting concentrated virus was used for infections as described in the “Viral Transduction” section. VZV titer was determined by digital droplet PCR (ddPCR) using primer/probes targeting ORF28 gene (Bio-Rad, Unique Assay ID: dCNS524077452).

### Cell protein extraction and digestion

As we previously reported, we used an automated sample preparation pipeline for protein extraction and digestion from cells for MS-based proteomics analysis ^36^. Briefly, the stock 4x lysis buffer consists of 200 mM Tris-HCI (pH 8.0), 200 mM NaCl, 1% SDS, 4% Triton X-100, 4% NP-40, 4% Tween 20, 4% glycerol, 4% Sodium deoxycholate (wt/vol), 20mM EDTA (pH 8.0). Before cell harvest, a working 1x lysis lysis buffer was prepared with final concentration of 5mM DTT, 5,000 units of benzonase, and 1× complete protease inhibitor. Cells were harvested from each well of a 6-well plate by gently washing with 500 µL of pre-chilled PBS buffer three times, being cautious to avoid lifting the cells off the plate. Then, 200 µL of the lysis buffer was added to each well, and the cells were scraped and transferred to 1.5 mL Eppendorf Safe-Lock tubes. Next, protein reduction and alkylation were performed by heating the cell lysate in a ThermoMixer at 65°C and 1200 rpm for 30 minutes. After the heating step, the tubes were removed and allowed to cool to room temperature. To alkylate the proteins, 4 µL of a 500 mM iodoacetamide stock solution was added to the lysate, bringing the final concentration of iodoacetamide to 10 mM. The lysate is incubated at room temperature in the dark for 30 minutes, with the tubes being wrapped in aluminum foil or placed in a dark room to prevent exposure to light. After the incubation, we normalized the protein concentration to 1mg/mL on a Bravo robot, and 40 µg proteins of each sample were transferred to a 96-deep well plate and enriched on a KingFisher robot using magnetic carboxylate modified particles. We digested the proteins using trypsin/LysC mix at 1:20 ratio on a ThermoMixer at 37°C and 1200 rpm for 16 hours. The resulting tryptic peptides were freezing dried and reconstituted in a 0.1% TFA, 2% ACN MS loading buffer.

### Liquid chromatography and mass spectrometry analysis

For peptides separation on a liquid chromatography (LC), 1 µg of the digested peptides were separated on a nano LC column, (75 μm × 500 mm, 2 μm C_18_ particle) using a 80min efficient linear gradient 2% - 35% phase B (5% DMSO in 0.1% formic acid, in Acetonitrile) with constant flow rate of 300 nL/min on an UltiMate 3000 nano-HPLC system. For the MS analysis, we used data independent acquisition (DIA) analysis where MS1 resolution was set to 120K, and both standard AGC target and auto maximum injection time were selected. For MS2 scans, the precursor range was set to 400-1000 m/z, and the isolation window was 8 m/z with 1 m/z overlap, resulting in 75 windows for each scan cycle. The MS2 fragmentation was performed in Orbitrap with 30K resolution and HCD with 30% collision energy, and the MS2 scan range was defined as 145-1450 m/z, and the loop control was set to 3 s.

### Proteomics database search

For protein annotation and quantification, we used the DIA-NN database search engine (version 1.9.2) to assign DIA spectra to peptides ^34^. A customized FASTA database was generated containing 20,406 human proteins and 365 viral proteins from the UniProt Consortium. Human and viral protein database consists of UP000005640 (*Homo sapiens*), UP000121444 (*HSV-1*), UP000006716 (*HCoV-229E*), UP000153037 (*EBV*), UP000002602 (*VZV*), UP000009255 (*H1N1*). DIA-NN was run using this customized FASTA database in the library free mode with Match Between Runs enabled. The database search considered 2 missed cleavage sites, methionine oxidation, and N-terminal acetylation. The results were filtered by a false discovery rate (FDR) of 1% on both precursor and protein level (Q value <0.01). Database searching utilized the computational resources of the NIH HPC Biowulf cluster (https://hpc.nih.gov).

### Post-analysis for proteomics data

Differential protein abundance analysis, functional enrichment, UMAP clustering, and visualization were performed using our in-house built proteomics informatic pipeline, ProtPipe ^40^. Briefly, proteins were considered differentially expressed across the comparisons if the absolute median ratio of two conditions was greater than 1.5 with an adjusted p-value < 0.05 (Benjamini– Hochberg adjusted p-value of a t-test). Functional enrichment analysis used adjusted p-value < 0.05 or q-value < 0.05 as the cut-off.

### WGCNA

We used the WGCNA R package^41^ (version 1.73) to perform protein network analysis. First, outlier samples and proteins from our data matrix were removed using the WGCNA::goodSamplesGenes() function. This removed samples and proteins with many missing values, weights below a threshold, or zero-variance variables. Additional outlier samples detected through PCA clustering were removed as well. The data was then divided into a dataset containing only human proteins and a dataset containing only viral proteins. Protein intensity values for the human dataset were log2 transformed and normalized by mean. The final matrix containing 145 sample rows and 8,929 protein columns was used in the WGCNA::blockwiseModules() function for network construction and protein module clustering.

We used a signed network type, maxBlockSize of 14000, mergeCutHeight of 0.25 and power of 12. The power parameter was specifically chosen as the minimum value needed to achieve scale free topology for our dataset. Using the viral protein data, we calculated the mean log2 protein intensity for each of the 5 viruses. We then determined the Pearson correlation coefficient between these aggregated viral protein values and the module eigengene for each module. Module eigengenes are defined as the first principal component of the protein expression values within each module. The correlation coefficients and associated p-values were used to determine the relationships between the 18 protein modules and the 5 viral infections.

### Cross validation with human cohort studies

To validate our findings, we utilized data from human cohort studies, which are detailed in Supplementary Table 2 (AD brain) and Supplementary Table 4 (AD CSF) of the publication ^17^. These studies included mass spectrometry (MS)-based proteomics data of postmortem brain tissues and CSF of AD patients and healthy controls. To identify differentially expressed proteins (DEPs) in the human cohort, we applied the following cutoff criteria: an absolute value of log2 FC ≥ 0.585 and an adjusted p-value < 0.05. Using these criteria, we performed an overlap analysis to identify proteins that were differentially expressed in both AD patient samples and viral-infected neurons. For the protein-protein interaction (PPI) analysis, we used the STRING database to generate and visualize the interactions between the identified DEPs.

## Supplementary Figure legends

**Figure S1. Quality Control of the Data**

(A) Number of human proteins identified across all samples.

(B) Uniform Manifold Approximation and Projection (UMAP) analysis showing clustering patterns of proteomic data based on the conditions of viral infection, suggesting the clusters of proteome profile is separated by different viral exposure.

**Figure S2. WGCNA Data Analysis**

(A) Plot showing the model fitting by the power soft threshold for network construction in WGCNA.

(B) Number of proteins identified in each module.

(C) Heatmap showing the log2 FC of neurodegenerative disease (NDD) genes across different viral infection conditions.

(D) PPI network of the genes identified in panel C, illustrating the interactions among dysregulated proteins involved in NDDs.

**Figure S3. Dysregulated Proteins Across All Comparisons**

(A) Heatmap showing the differential expression of genes across all virus types and time points, highlighting the proteins most significantly dysregulated.

(B) UpSet plot identifying genes commonly associated with infection across all viruses, illustrating overlaps in gene dysregulation between the different viral infections.

**Figure S4. WGCNA Module GO Pathway Analysis**

(A) Module 16 GO pathway analysis related to wound healing, showing the significantly enriched biological processes and pathways involved in wound repair and tissue regeneration.

(B) Module 10 KEGG pathway analysis related to protein degradation, highlighting pathways such as lysosome and ubiquitin-proteasome systems that are crucial for cellular protein turnover.

(C) Module 5 KEGG pathway analysis for DNA repair and cell cycle processes, illustrating key pathways involved in maintaining cellular integrity and function.

(D) Module 2 analysis focused on mRNA splicing and processing, showing the pathways related to RNA metabolism and regulation.

## Supplemental tables

Supplemental table1 human protein count

Supplemental table2 viral protein count

Supplemental table3 DEP results of virus-infected neurons compared to controls

Supplemental table4 DEP pathway enrichment results of virus-infected neurons compared to controls

Supplemental table5 WGCNA module membership

