## Supplementary figures and images for "Proteomic Analysis of Endemic Viral Infections in Neurons offers Insights into Neurodegenerative Diseases"

### Supplemental Figures

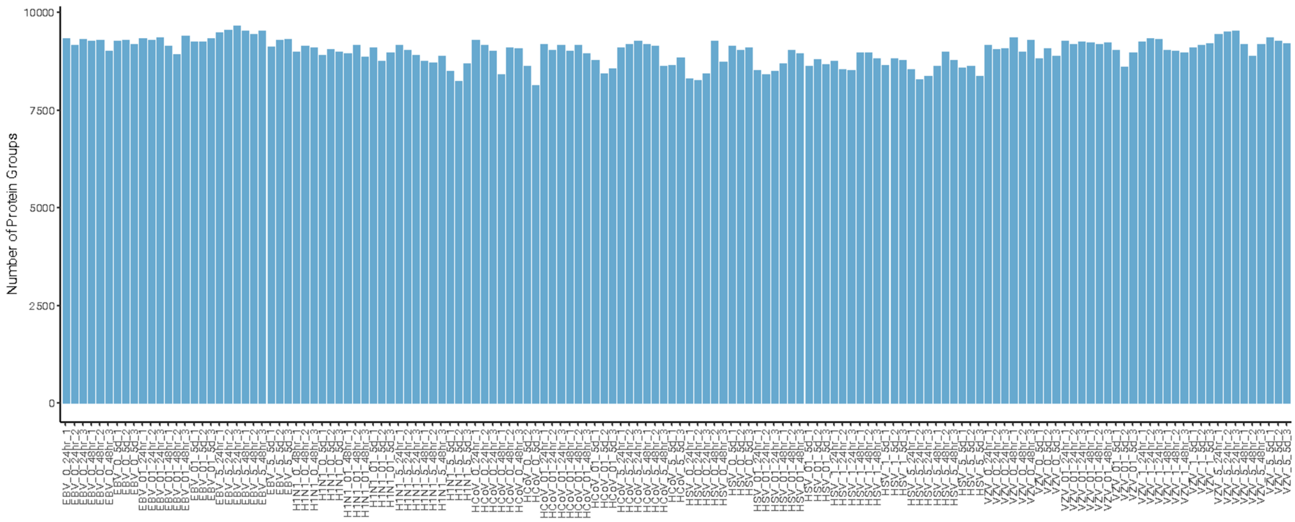

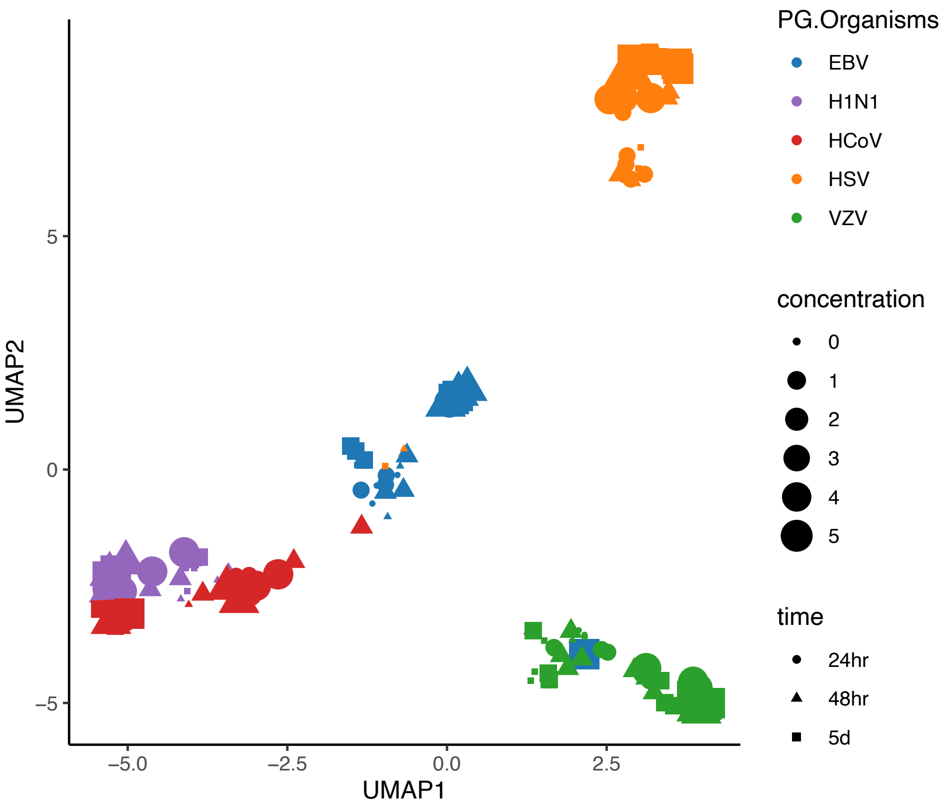

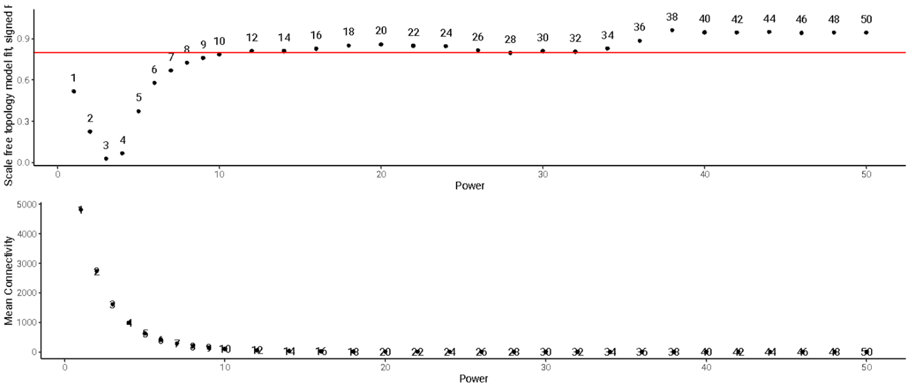

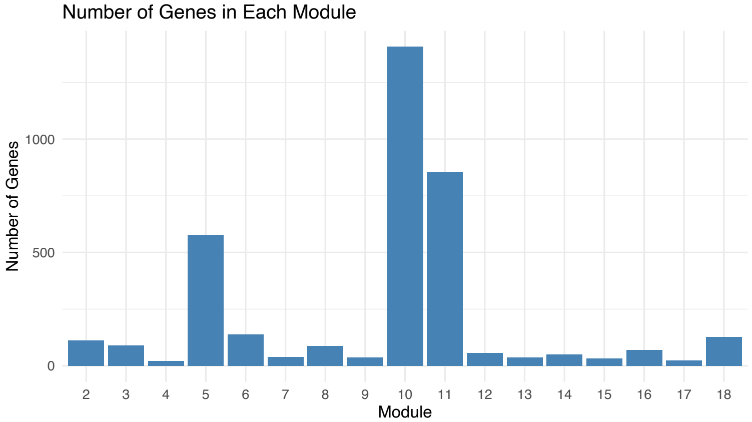

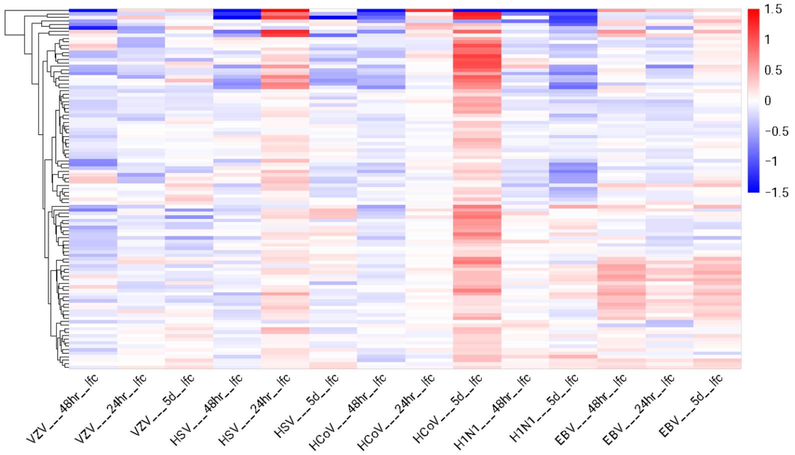

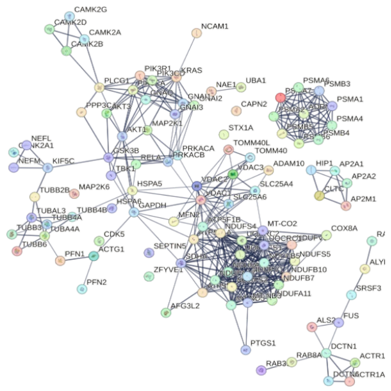

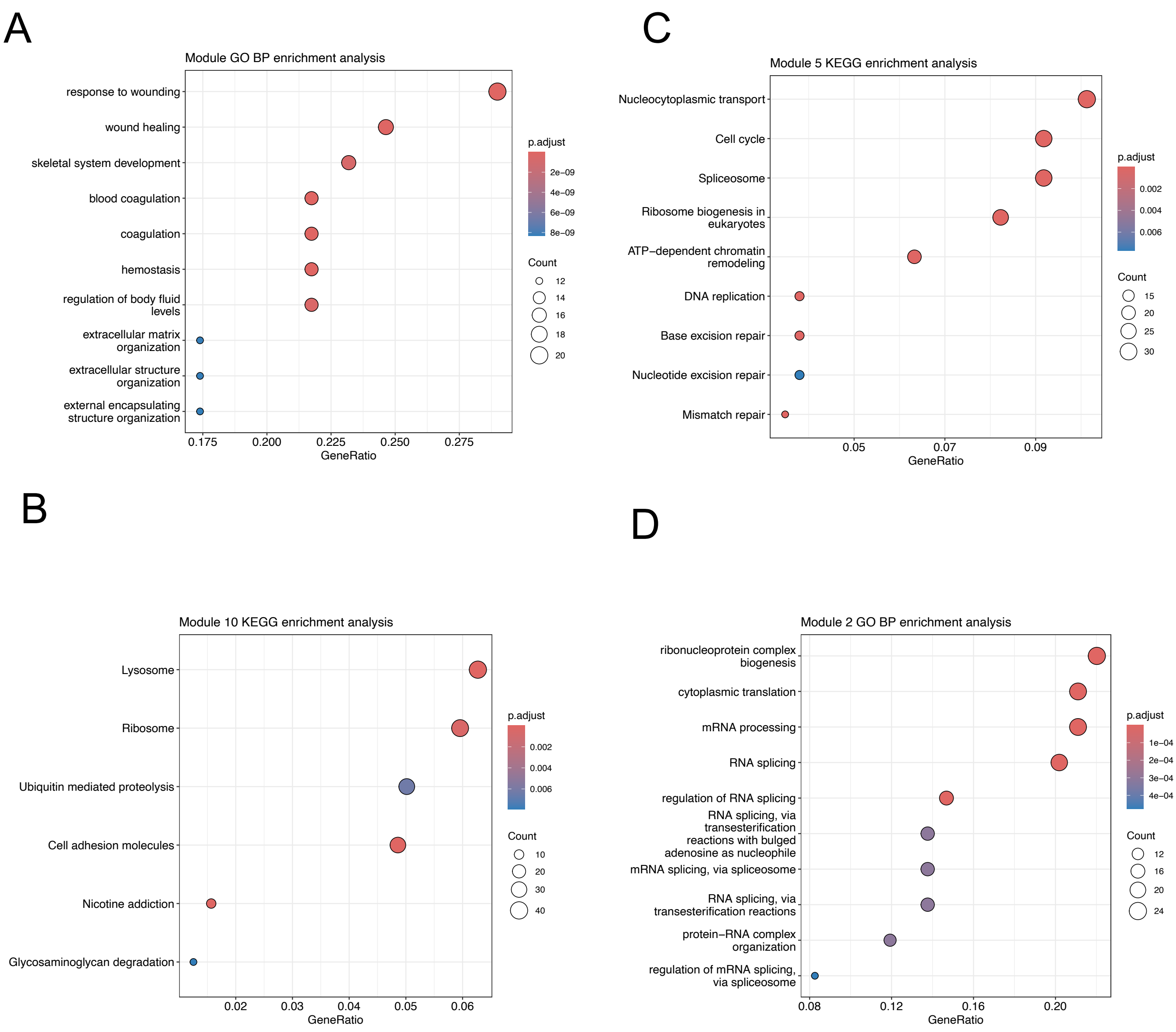
